# Atypical MAPK regulates translocation of GATA transcription factor in response to chemoattractant stimulation

**DOI:** 10.1101/2022.02.17.480945

**Authors:** Jeffrey A. Hadwiger, Huaqing Cai, Ramee G. Aranda, Saher Fatima

## Abstract

The *Dictyostelium* atypical MAPK Erk2 (a.k.a. ErkB) is required for chemotactic responses to external cAMP as individual amoeba aggregate and undergo a multicellular developmental program. External cAMP also stimulates the translocation of the GATA transcription factor, GtaC, a key regulator of developmental gene expression and a developmental timer of repeated cAMP stimulation of developing cells. In this study, atypical MAPK Erk2 was found to be essential for the translocation of GtaC in response to external cAMP as indicated by the cellular distribution of GFP-GtaC reporters. Erk2 was also demonstrated to mediate the translocation of GtaC in response to external folate, a signal that directs the foraging of amoeba for bacterial food sources. Erk1, the only other *Dictyostelium* MAPK, was not required for the GtaC translocation to either chemoattractant, indicating that GFP-GtaC is a kinase translocation reporter specific for atypical MAPKs. The translocation of GFP-GtaC in response to folate was absent in mutants lacking the folate receptor, Far1 (a.k.a. GrlL) or the coupled G protein, Gα4 subunit. Loss of GtaC function resulted in enhanced chemotactic movement to folate. Inspection of the GtaC primary sequence revealed four sites recognized as Erk2 preferred phosphorylation sites. The creation and analysis of GFP-GtaC mutants with alterations at these sites determined that these sites are required for translocation in response to folate. Consistent with the role of translocation for GtaC function certain combinations of these phosphorylation site alterations interfered with the ability of GFP-GtaC reporter to rescue aggregation of *gtaC^-^* cells. These findings provide the first evidence that atypical MAPKs can regulate transcription factors through specific phosphorylation sites in response to chemoattractants. The ability of different stimuli to regulate GtaC translocation through Erk2 indicates atypical MAPKs can regulate different cell fates in part through the translocation of transcription factors.

## Introduction

Environmental signals can regulate eukaryotic cell function and fate through signaling pathways that control cellular processes such as gene expression and cell movement. These signaling pathways typically involve cell surface receptors coupled to G proteins and multiple protein kinases to serve as molecular switches for a wide range of cellular responses (Cargnello and Roux, 2011; Goldsmith and Dhanasekaran, 2007; Luttrell, 2006). Mitogen activated protein kinases (MAPKs) play important roles in many different signaling pathways in most eukaryotes, including animals, plants, fungi, and protists (Bogoyevitch and Court, 2004; Chen and Thorner, 2007; Doczi et al., 2012; Hadwiger and Nguyen, 2011; Yildiz and Arslanyolu, 2014). Theses kinases are typically downstream of other protein kinases in G protein-coupled receptor and tyrosine kinase receptor signaling pathways and they facilitate the regulation of many types of signaling proteins including other protein kinases, phosphodiesterases, and transcription factors that modulate cellular activities such as second messenger signaling, metabolism, chemotaxis, and gene expression (Cargnello and Roux, 2011; MacKenzie et al., 2000; Maeda et al., 2004; Nichols et al., 2019; Roskoski, 2012; Schwebs et al., 2018). Canonical MAPK regulation involves kinase cascades that include MAPK kinases (MAP2K), MAPKK kinases (MAP3K), and scaffolding proteins. However, one subgroup of MAPKs referred to as atypical MAPKs do not appear to be regulated through conventional MAP2Ks but instead have another regulatory mechanism that remains to be fully understood (Coulombe and Meloche, 2007). Compared to other MAPKs, relatively little is known about the function of atypical MAPKs in animals but several reports have indirectly associated atypical MAPKs with stress responses (Chia et al., 2014; Colecchia et al., 2018; Groehler and Lannigan, 2010; Hasygar and Hietakangas, 2014; Iavarone et al., 2006; Klevernic et al., 2009; Liwak-Muir et al., 2016; Rossi et al., 2011; Saelzler et al., 2006; Zacharogianni et al., 2011). Challenges to understanding the role of atypical MAPKs are identifying specific signals that directly stimulate atypical MAPK activation and characterizing the phenotypes that result from losses of atypical MAPK function, but the use of genetically amenable organisms has begun to bring much insight into the roles of these protein kinases in cell fate regulation (Chen and Segall, 2006; Gaskins et al., 1996; Maeda et al., 1996; Maeda and Firtel, 1997; Schwebs et al., 2018; Segall et al., 1995).

Atypical MAPKs have been found in animals and protists, such as amoebae, and they share some structural similarities that make them distinct from other MAPKs (Abe et al., 2002; Ellis et al., 2004; Segall et al., 1995; Valenciano et al., 2016). The absence of atypical MAPKs in fungi has excluded their analysis in yeast which have provided many insights into function and regulation of other types of MAPKs (Chen and Thorner, 2007). However, the amoeba *Dictyostelium discoideum* encodes an atypical MAPK and genetic analyses have provide many clues to atypical MAPK function and regulation (Schwebs et al., 2018). The *Dictyostelium* atypical MAPK, Erk2, is rapidly activated (within 30 seconds) in response to chemoattractants and mediates chemotactic cell movement, consistent with presence of atypical MAPKs in organisms with cell motility (Maeda et al., 1996; Maeda and Firtel, 1997; Nichols et al., 2019; Schwebs et al., 2018). *Dictyostelium* encodes only one other MAPK, Erk1, that is related to other more typical MAPKs found in most all eukaryotes (Gaskins et al., 1994; Schwebs et al., 2018). Both Erk2 and Erk1 can become activated through the phosphorylation of an activation motif (TEY) in response to chemotactic signaling but with very different kinetics and activators (Schwebs and Hadwiger, 2015). Erk2 activation occurs first and then Erk1 activation occurs after a few minutes as Erk2 becomes dephosphorylated. The single conventional MAP2K in *Dictyostelium* is only required for the phosphorylation of Erk1 and not Erk2 (Schwebs et al., 2018). Erk2 is the only MAPK required for chemotactic movement.

*Dictyostelium* chemotax to at least two chemoattractants, cAMP and folate (Gerisch, 1982). Chemotaxis to folate is a mechanism by which *Dictyostelium* can forage for bacterial food sources in the environment and resume vegetative growth as solitary cells. In contrast, chemotaxis to cAMP mediates the aggregation phase of the multicellular developmental life cycle by allowing cells to move toward each other as they relay cAMP signaling in a pulsatile manner. Responses to cAMP not only include chemotactic movement but the regulation of many developmental genes, including developmental cell type specific genes (Rosengarten et al., 2015). One of the key regulators of developmental genes is the GATA transcription factor GtaC (Cai et al., 2014; Keller and Thompson, 2008; Santhanam et al., 2015). Loss of GtaC impairs aggregation and GtaC has been shown to bind at many locations in the genome (Santhanam et al., 2015). GFP-tagged GtaC rescues the development of *gatC*^-^ strains and reports the translocation of this transcription factor from the nucleus to the cytoplasm when cells are stimulated with cAMP (Cai et al., 2014). The shuttling of this transcription between the nucleus and cytoplasm during the repeated cyclic pattern of cAMP stimulation has been suggested to function as a developmental timer in the assessment of cycle number and duration. In this report we assessed the roles of both typical and atypical MAPKs in the translocation of this GATA transcription factor in response to multiple chemoattractant signals and also tested the specificity of chemoattractant receptors and associated G proteins in this process. The GFP-GtaC reporter was used to examine the roles of specific phosphorylation sites on GtaC translocation and the regulation of developmental processes. Our study suggests that multiple external signals can use atypical MAPK signaling to regulate this transcription factor and different cell fates.

## Results

### Atypical MAPK Erk2 is required for GtaC translocation

The translocation of a GFP-GtaC reporter from the nucleus to the cytoplasm was previously observed in a strain containing a hypomorphic *erk2*^-^ allele but we reexamined the possible translocation of the reporter in an *erk2*^-^ null strain (Cai et al., 2014). Transformants containing the GFP-GtaC reporter (pHC326) were only obtained with a low concentration drug selection (1.5 μg G418/ml) and none of the transformants displayed sufficient GFP fluorescence for analysis, suggesting that the reporter might be toxic to cells lacking Erk2. To reduce the possible toxicity of GFP-GtaC reporter a mutant version of the reporter, GFP-GtaC^c-s^ (pHC329), was used in which two of the required cysteine residues of the GtaC zinc finger domain were converted to serine residues (C500S/C503S). Many viable transformants were obtained using this reporter vector and GFP fluorescence was readily detectable at high drug selections (3-8 μg/ml) suggesting the alteration of the zinc finger reduces toxicity of the reporter. The reporter was concentrated in the nucleus of the *erk2* null cells during both growth and starvation and the stimulation of cells with cAMP did not result in the translocation of the reporter to the cytoplasm (Fig. 1, video S1). Complementation of the *erk2* null mutant with a wildtype *erk2* allele rescued the ability of cAMP stimulation to translocate the reporter to the cytoplasm confirming that Erk2 is essential for this process. The translocation of the reporter was also re-examined in an *erk1* null strain and was found to translocate comparable to wild-type cells confirming that Erk1 function is not required for this process.

**Figure 1.**
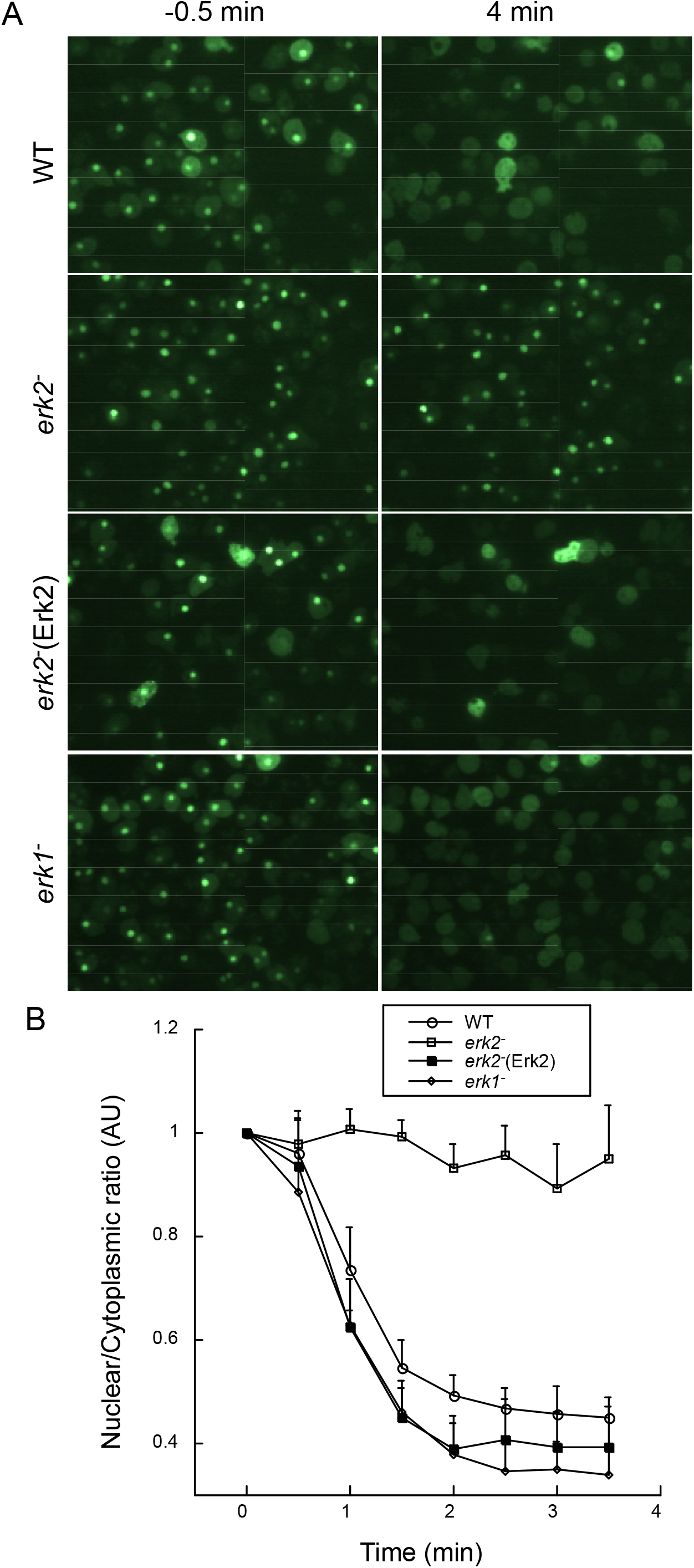
Requirement of Erk2 function for the translocation of GtaC in response to cAMP stimulation. (A) Wild-type (WT), *erk2*^-^, *erk2*^-^ complemented with Erk2 expression vector (Erk2), and *erk1*^-^ strains expressing the GFP-GtaC^c-s^ reporter were stimulated with 10 nM cAMP and time-lapse images were collected every 30 sec. Images of −0.5 and 4 min are displayed of representative assay of at least 3 independent assays. All images are the same magnification. (B) Graphical representation of mean nuclear/cytoplasmic ratios (arbitrary units) of WT (n=35), *erk2^-^* (n=20), *erk2^-^* (ERK2) (n=35), and *erk1^-^* (n=40) strains. Error bars represent + standard deviation.

### Erk2 is required for phosphorylation of GtaC

The stimulation of wild-type cells with cAMP results in the rapid appearance of phosphorylated forms of the GFP-GtaC reporter that migrate slower in electrophoretic gels and an analysis of GtaC sequence revealed many potential phosphorylation sites of several types of protein kinases (Cai et al., 2014). The slower migrating forms of GFP-GtaC were absent in *erk2^-^* cells stimulated with cAMP suggesting that Erk2 is required for the phosphorylation of the reporter (Fig. 2). The slower migrating forms of the reporter were observed in *erk2-* cells complemented with the wild-type *erk2* gene and also in *erk1^-^* cells, suggesting a specific dependence on Erk2 function. The requirement of Erk2 function for GtaC phosphorylation does not exclude the possibility that other protein kinases, such as glycogen synthase protein kinase (gsk3), might phosphorylate GtaC but only the loss of Erk2 prevents the mobility shift of the GFP-GtaC reporter (Cai et al., 2014). In an *in vitro* phosphorylation assay immunoprecipitated Erk2, but not Erk1, was capable of phosphorylating GFP-GtaC resulting in a mobility shift in in electrophoretic gels (Fig. 2B). In total, these data suggest GtaC is specifically and directly phosphorylated by Erk2, a mechanism consistent with the dependence of GtaC translocation on Erk2 function.

**Figure 2.**
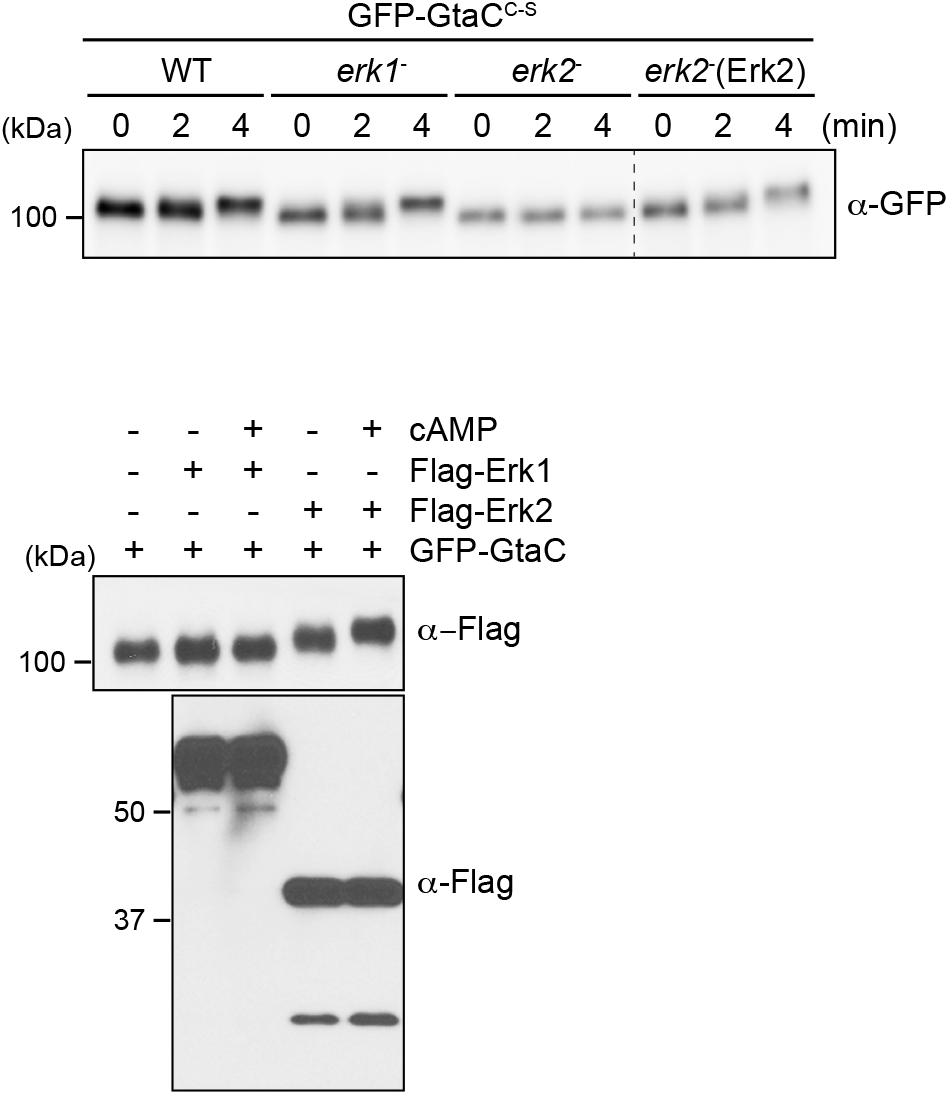
Requirement of Erk2 function for GFP-GtaC mobility shifts. (A) Immunoblot of GFP-GtaC^c-s^ from wild-type (WT), *erk1^-^, erk2^-^*, and *erk2^-^* complemented with Erk2 expression vector (Erk2) strains stimulated with cAMP. Dashed line indicates removal of extra lanes in blot. (B) GFP-GtaC purified from wild-type cells and Flag-Erk1 and Flag-Erk2 purified from wild-type cells with or without cAMP treatment were combined in an *in vitro* kinase assay. Immunoblot of GFP-GtaC^c-s^ mobility shifts and immunoblots of Flag-Erk1 (~ 70 kDa) and Flag-Erk2 (~42 kDa).

### Erk2 regulates GtaC translocation in response to folate

Erk2 is activated in response to at least two chemoattractants, cAMP and folate, and we postulated that the activation of Erk2 in response to folate could possibly also lead to GtaC regulation. Although GtaC translocation and function have been associated with cAMP signaling and gene expression, it might also be important for other responses to nutrient limitation. Folate stimulation of wild-type cells resulted in the translocation of the GFP-GtaC^c-s^ reporter from the nucleus to the cytoplasm with kinetics comparable to that occurring after cAMP stimulation (Fig. 3, video S2). This translocation was absent in *erk2*^-^ cells similar to that observed for cAMP stimulated cells. The translocation of the reporter was rescued by the expression of wild-type Erk2 indicating that Erk2 can regulate GtaC translocation in response to multiple chemotactic signals. As with cAMP responses, Erk1 was not required for the translocation of the reporter in response to folate implying that only atypical MAPK function regulates this process.

**Figure 3.**
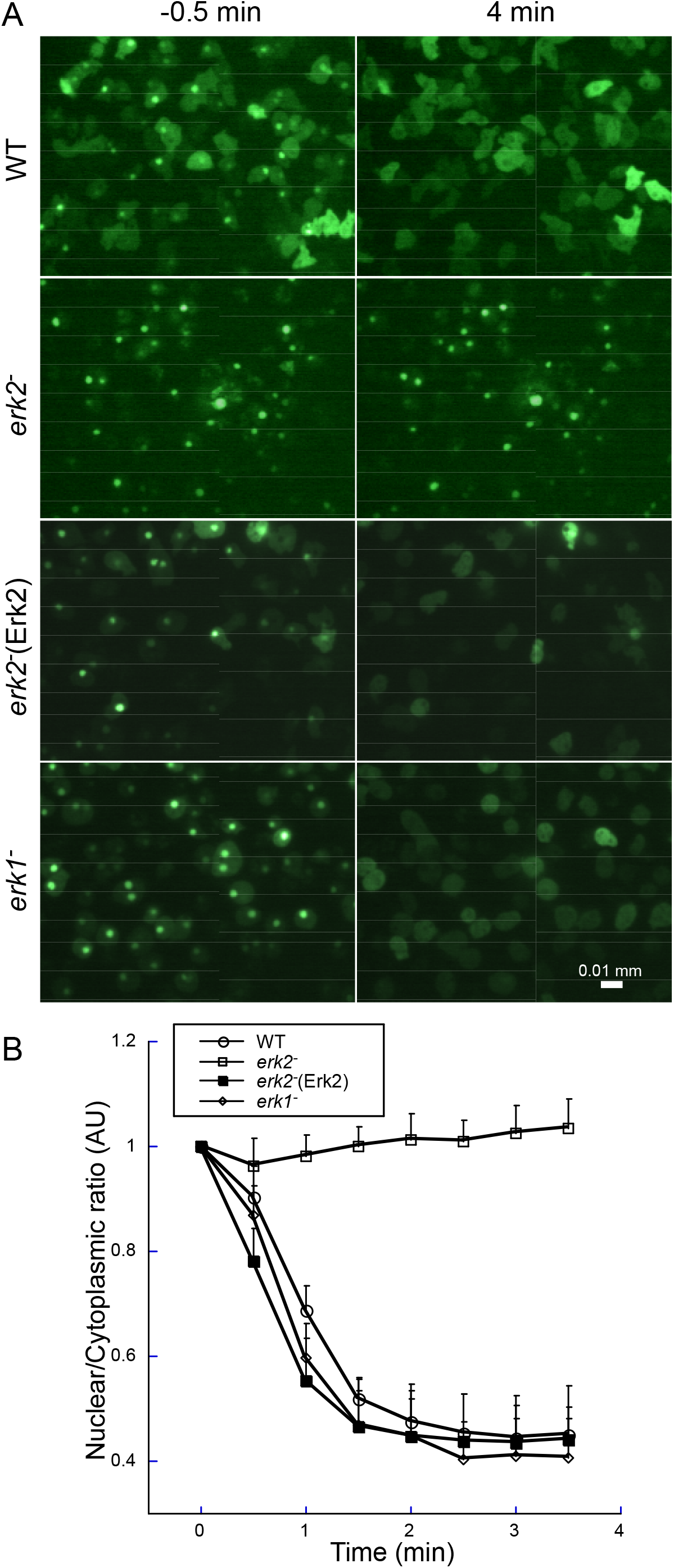
Erk2 dependency of the GtaC translocation in response to folate. (A) Wildtype (WT), *erk2*^-^, *erk2*^-^ complemented with Erk2 expression vector (Erk2), and *erk1*^-^ strains expressing the GFP-GtaC^c-s^ reporter were stimulated with 1 μM folate and time-lapse images were collected every 30 sec as described in Fig. 1. Images of −0.5 and 4 min are displayed of representative assay of at least 3 independent assays. All images are the same magnification. (B) Graphical representation of mean nuclear/cytoplasmic ratios (arbitrary units) of WT (n=23), *erk2*^-^ (n=31), *erk2*^-^(ERK2) (n=25), and *erk1*^-^ (n=40) strains. Error bars represent + standard deviation.

### Folate signaling and GtaC function

Chemotaxis and other responses to folate require the folate receptor, Far1, and the coupled G protein subunit, Gα4, and so the requirements of these signaling components were analyzed by assaying the GFP-GtaC^c-s^ reporter translocation in cells lacking either the receptor or the G protein. As expected, folate stimulation of the reporter translocation was absent in cells lacking the folate receptor Far1 and in cells lacking G protein subunit Gα4 (Fig. 4, video S3). The apparent dip in the nuclear/cytoplasmic ratio of GFP-GtaC in *gα4^-^* cells stimulated with folate was observed in multiple assays and appeared to result from a transient change in shape of some cells that temporarily moved the nucleus out of the focal plane. The loss of either Far1 or Gα4 did not prevent translocation of the reporter when cells were stimulated with cAMP, confirming that these signaling proteins mediate responses to folate and not cAMP. The closest Gα4 G protein paralog in *Dictyostelium* is the Gα5 subunit and loss of the Gα5 subunit did not impede translocation of the reporter in response to either folate or cAMP. Previous studies have shown the Gα5 subunit is not required for chemotactic movement to folate but possibly acts in opposition to the Ga4 subunit in chemotactic movement and developmental morphology (Natarajan et al., 2000). Earlier studies have also indicated the requirement of cAMP receptors, Car1 and Car3, and the G protein subunit Gα2 for efficient translocation of the reporter in response to cAMP (Adhikari et al., 2021; Cai et al., 2014). To evaluate GtaC function on folate chemotaxis, a *gtaC^-^* strain was created through gene disruption and analyzed for chemotaxis to folate in above agar assays. Cells lacking GtaC migrated further toward folate than the wild-type parental cells consistent with a role for GtaC in the suppression of Far1 expression (Fig. 4C, S4). This result suggests that while Erk2 is required for folate chemotaxis the Erk2 mediated translocation of GtaC might enhance chemotaxis through increased expression of the Far1 receptor.

**Figure 4.**
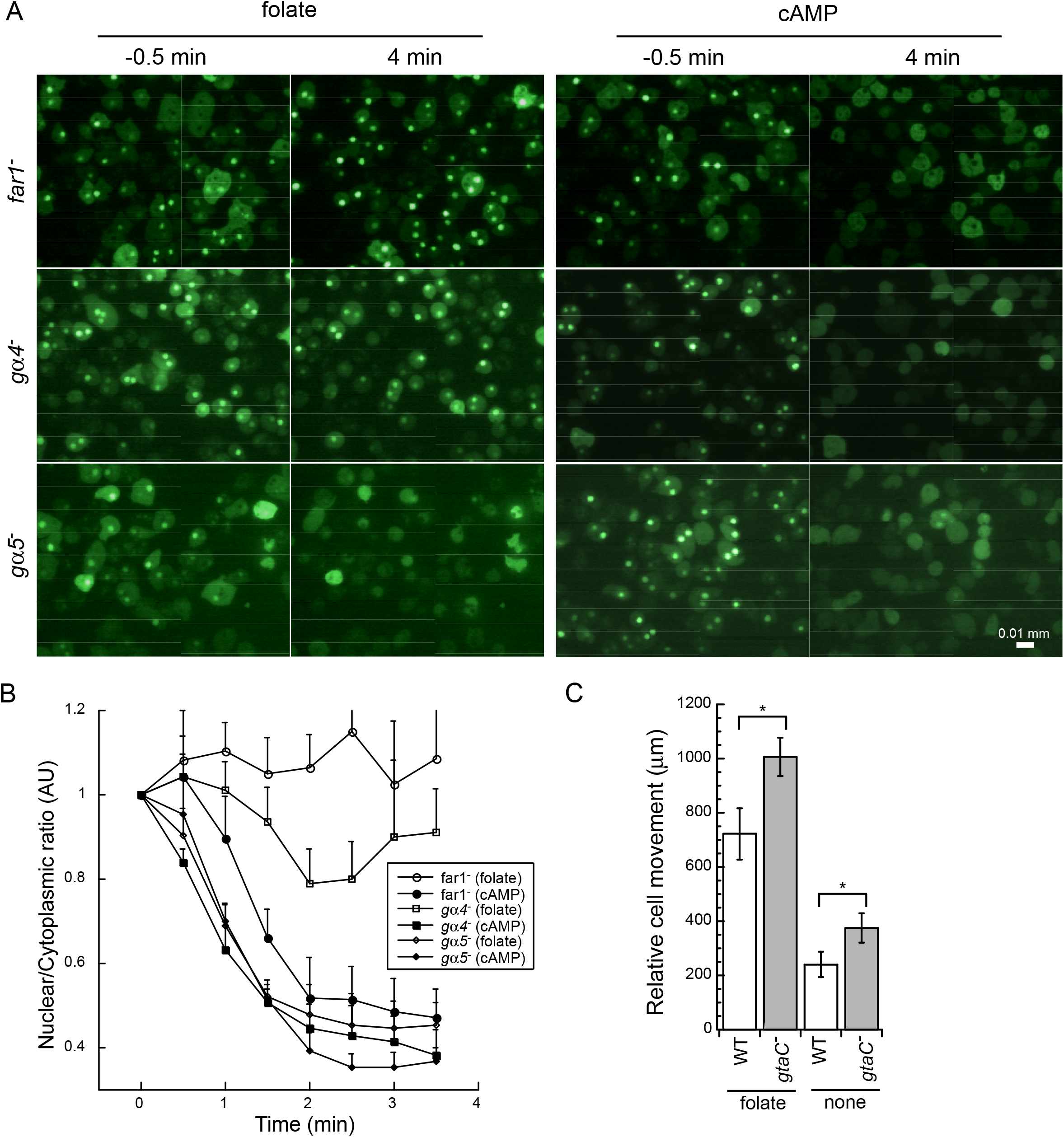
Receptor and G protein requirements for GtaC translocation in response to folate. (A) far1 -, ga4-, and ga5-strains expressing the GFP-GtaC^c-s^ reporter were stimulated with 1 μM folate or 10 nM cAMP and time-lapse images were collected every 30 sec as described in Fig. 1. Images of −0.5 and 4 min are displayed of representative assay of at least 3 independent assays. See supplemental Fig. X for videos of the timelapse images. All images are the same magnification. (B) Graphical representation of mean nuclear/cytoplasmic ratios (arbitrary units) of far1 (folate n=28, cAMP n=30), *ga4*^-^ (folate n=30, cAMP n=40), *gα5*^-^ (folate n=23, cAMP n=25) strains. Error bars represent + standard deviation. (C) Above agar chemotaxis assays of wild-type (WT) and *gtaC^-^*cells to folate. Data shown is representative of at least four assays. Error bars represent standard deviation of means. * indicates student’s t-test p < 0.001.

### Erk2-dependent translocation of GtaC can be mediated through multiple phosphorylation sites

The dependence of GtaC phosphorylation and translocation on Erk2 function suggests that Erk2 phosphorylates GtaC before it can be translocated to the cytoplasm. An earlier analysis of Erk2 phosphorylation sites in vivo and in vitro has indicated that preferred Erk2 phosphorylation sites include serine/threonine residues followed by a proline and a positively charged residue (arginine/lysine) (Nichols et al., 2019). The primary sequence of GtaC contains four of these preferred sites (S357, S380, S386, and T492) whereas none of these sites exist in the GFP tag. Of these four sites, only the T492 residue is located within a region highly conserved among other *Dictyostelid* GtaC homologs and other sites exist just upstream of this conserved region (Fig. S5). Other GtaC orthologs also have putative Erk2 phosphorylation sites just upstream of the conserved region but the positions of these sites do not always align precisely with those in the *Dictyostelium discoideum* GtaC protein. In addition, GtaC contains 12 other sites (S/T-P) that are generally regarded potential targets for MAPKs or cyclin-dependent kinases. An increase in the phosphorylation of the T492 site has been indicated in a mass spectrometry analysis of phosphoprotein after folate stimulation (Nichols, et al.). We examined the importance of these four putative Erk2 preferred phosphorylation sites for the translocation of the GFP-GtaC reporter in response to folate stimulation by converting one or more of the target residues (serine or threonine) to alanine. Individual residue changes did not prevent the translocation of GFP-GtaC^c-s^ reporter in wild-type cells but in some cases the extent of translocation was reduced (Fig. 5, video S6). These observations indicate that no single site is essential and that redundancy might exist in the ability of these sites to allow translocation of the reporter. Converting all of the sites or all but the S357 site significantly interfered with the reporter translocation and in general the reporter translocation decreased as more sites were altered. Conversion of S380 and/or S386 appeared to have more impact on reducing reporter translocation than other single or double conversions suggesting these residues might be the primary targets of Erk2 phosphorylation. However, the relationship of specific conversions with the respective impact on reporter translocation appears to be complex and the possibility cannot be ruled out that a conversion of one site might indirectly impact the importance of other sites.

**Figure 5.**
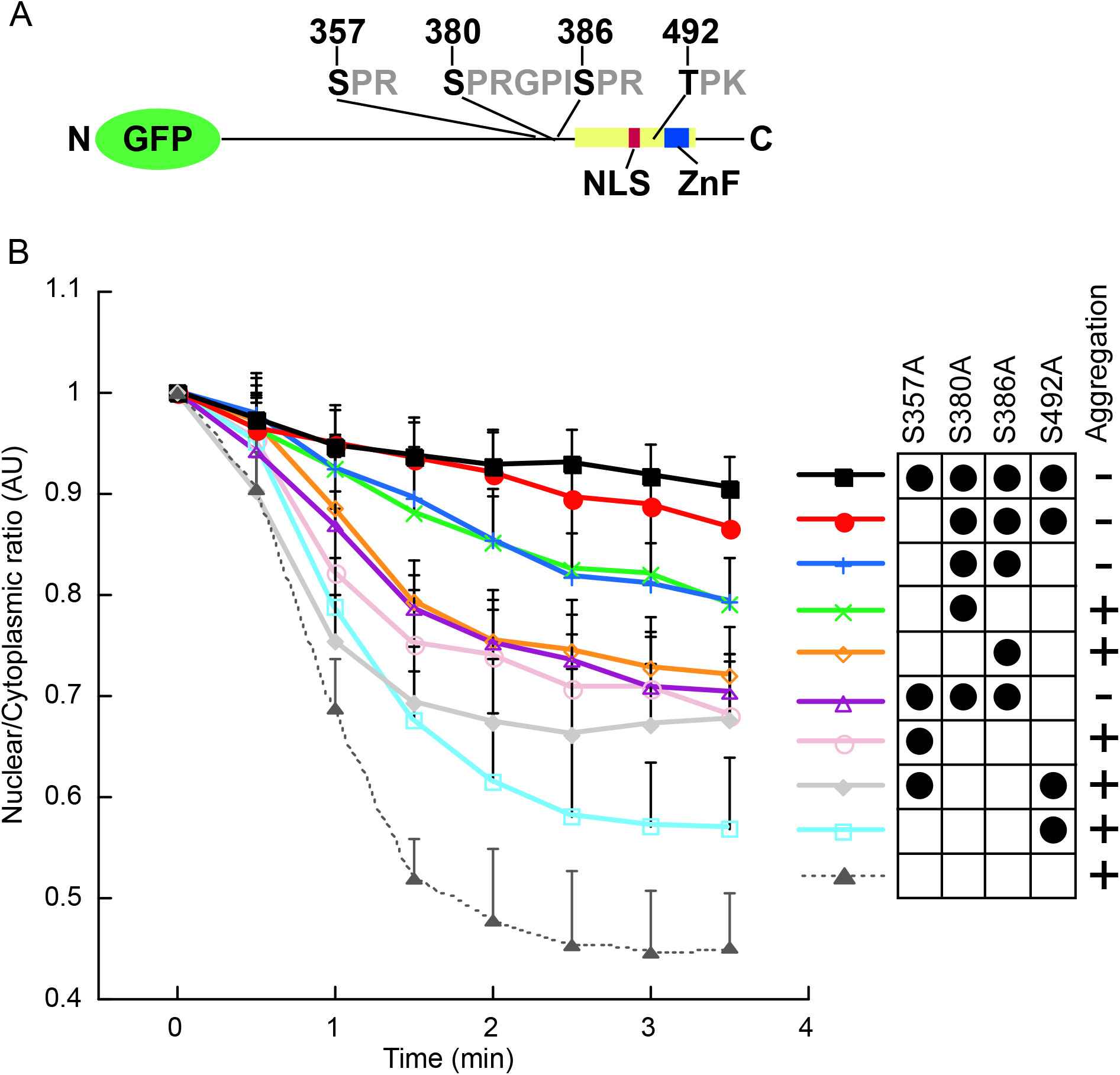
Erk2 phosphorylation requirements for GtaC translocation in response to folate. (A) Map of GFP-GtaC reporter construct with locations of Erk2 preferred phosphorylation motifs. Location of nuclear localization signal (NLS) (red bar) and zinc finger domain (ZnF) (blue bar) are indicated within highly conserved region (yellow bar, see also Fig. S6) of Dictyostelid homologs. (B) Graphical representation of nuclear/cytoplasmic ratios of GFP-GtaC^c-s^ mutants expressed in wild-type cells after stimulation with 1 μM folate. Time-lapse images were collected and analyzed as described in Fig. 1 and Methods. Wild-type cells with the parental vector (pHC329) data from Fig. 1 is included for comparison. Key indicates the combination of Erk2 phosphorylation site alterations, graph line color, and vector number. Multiple representative assays of wild-type cells with pJH911 (n=56), pJH913 (n=54), pJH925 (n=74), pJH926 (n=59), pJH929 (n=97), pJH931 (n=81), pJH932 (n=77), pJH935 (n=75), pJH937 (n=55), and pHC329 (n=23) were used to generate the mean of nuclear/cytoplasmic ratios (arbitrary units) and error bars indicated + standard deviations. (C) Aggregation capability of *gtaC^-^* cells expressing Erk2 phosphorylation mutant vectors with functional zinc finger domains (targeted sequence reversion) on bacterial lawns (images displayed in Fig. 6)

### Erk2 phosphorylation sites are necessary for GtaC function in early development

The GFP-GtaC^c-s^ reporter offers a real-time analysis of Erk2 mediated GtaC translocation but the translocation analysis might not accurately reflect the importance of Erk2 phosphorylation sites on GtaC function, particularly since the reporter lacks a functional zinc finger domain. To examine the impact of the Erk2 phosphorylation site alterations on GtaC function the zinc finger domain in these mutant alleles was restored by converting the zinc finger domain sequence back to the wild-type sequence (S500C/S503C). When introduced into a *gtaC^-^* strain all of the single phosphorylation site mutants and one of the double site mutants (S357A/T492A) were capable of rescuing the aggregation phase of development when cells were grown on bacterial lawns (Fig. 6). Interestingly, the S380A allele rescued aggregation to form tight mounds, but most mounds did not proceed further in the developmental progression. All of the combinational mutants possessed mutations at both S380 and S386 and none of these were capable of restoring aggregation necessary to form tight mounds of cells. These results indicate that Erk2 phosphorylation sites on GtaC are not just important for translocation but they also play a regulatory role in GtaC function consistent with a previous study that demonstrated a correlation between translocation and developmental gene expression (Cai et al., 2014).

**Figure 6.**
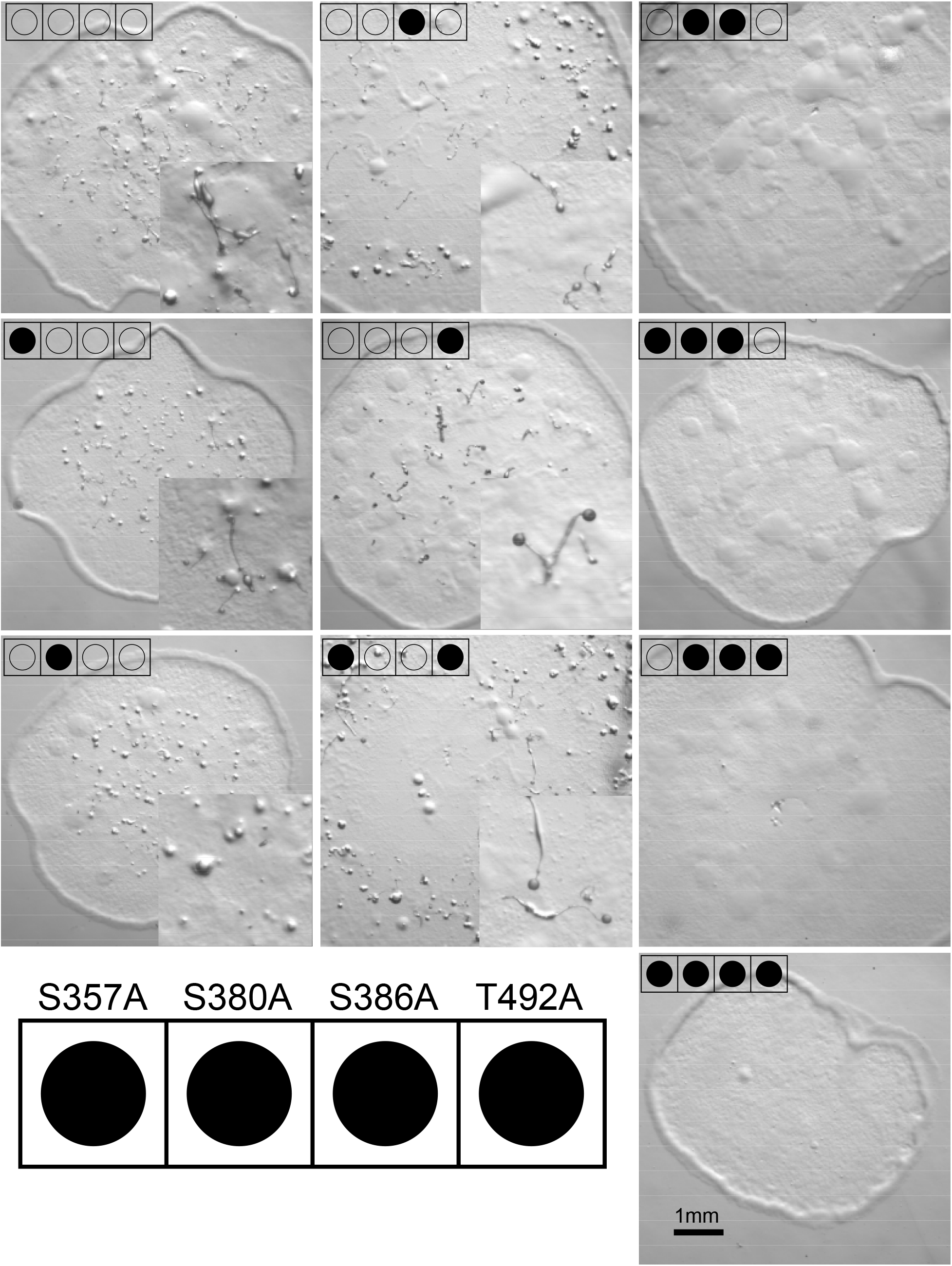
Erk2 phosphorylation site regulation of GtaC function in early development. GFP-GtaC constructs with altered Erk2 phosphorylation sites were expressed in *gtaC^-^* cells. Strains expressing each construct were plated on to bacterial lawns and allowed to grow and develop for 2-3 days. All images are the same magnification except for insets (3x magnification). Representative aggregates that have formed tight mound are indicated by arrows. Key indicates which sites have been altered (filled in circles) for each construct.

## Discussion

The dependence of GtaC translocation on the presence of the atypical MAPK Erk2 and on the Erk2 preferred phosphorylation sites within this transcription factor provide strong support that Erk2 directly regulates GtaC function in response to chemoattractants (Fig. 7). The interaction between Erk2 and GtaC is further supported by the phosphorylation of GtaC in combined immunoprecipates of GtaC and Erk2 as indicated by mobility shifts of GFP-GtaC in electrophoretic gels. These results reveal a previously unrealized role for atypical MAPKs in the regulation of transcription factor translocation suggesting that this class of MAPKs can directly modify key regulators of gene expression. This study demonstrates that Erk2 can phosphorylate and regulate the translocation of the GtaC transcription factor in response to multiple chemoattractants suggesting that the regulation of GtaC might have roles in foraging responses in addition to its role in multicellular development. The regulation of transcription factors in response to external signals has been observed for other classes of MAPKs. Mammalian ERK1/2 regulates the activation and nuclear accumulation of the transcription factor Elk-1 and p38 regulates the translocation of TEAD transcription factors from the nucleus to the cytoplasm in response to stress (Lavaur et al., 2007; Lin et al., 2017; Slone et al., 2016; Yang et al., 1999). Heterologous over-expression of the atypical MAPK, MAPK15, in mammalian cells has been reported to regulate CRM1-dependent ERRαlocalization to the cytoplasm through protein-protein interactions but the phosphorylation state of ERRa was not investigated (Rossi et al., 2011). Our analysis showing Erk2 regulation of GtaC in *Dictyostelium* is the first report of transcription factor phosphorylation and translocation by an atypical MAPK. It is important to note that the Erk2 regulation of GtaC is rapid and the timing correlates with the rapid activation of Erk2 in response to chemoattractants. Other classes of MAPKs, including Erk1 in *Dictyostelium,* typically display slower activation responses suggesting that the kinetics of atypical MAPK activation and function might be the primary MAPK response as observed in *Dictyostelium* chemotactic responses (Schwebs and Hadwiger, 2015).

**Figure 7.**
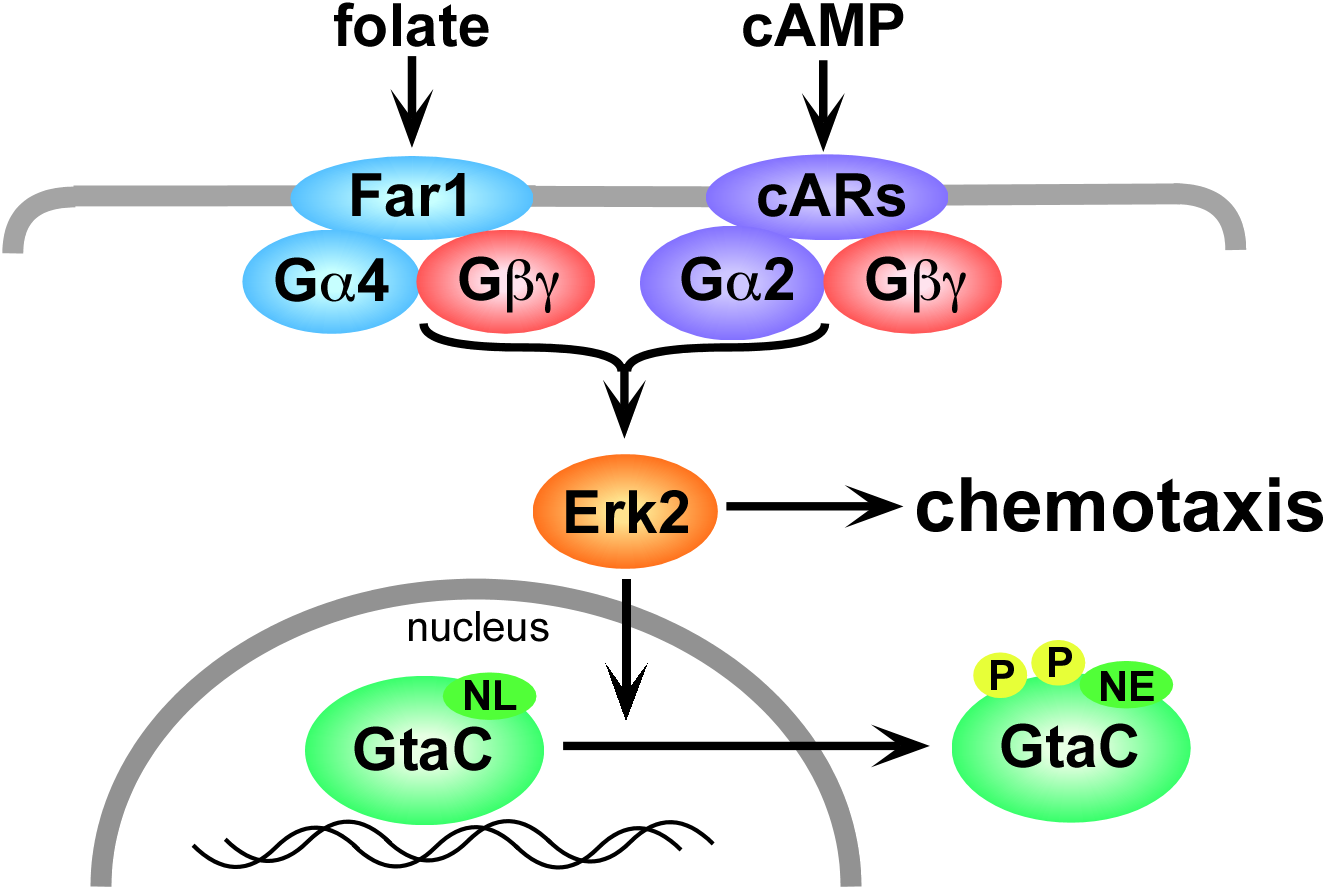
Model of Erk2 regulation of GtaC translocation. Activation of cAMP (cARs) or folate (Far1) G protein coupled receptors leads to the activation of the atypical MAPK Erk2 and the phosphorylation of the GtaC transcription factor. Translocation of GtaC from the nucleus to the cytoplasm is possibly mediated through masking the nuclear localization signal (NL) or exposing a nuclear export signal (NE).

Erk2 function is essential for folate chemotaxis and the loss of GtaC increases the movement of cells to folate suggesting Erk2 regulation of GtaC results in the increased expression of genes necessary for this response. An analysis of GtaC function on gene regulation has indicated that loss of GtaC function increases the expression of the folate receptor Far1 consistent with increased chemotactic movement to folate (Santhanam et al., 2015). Erk2 is likely to have other cellular targets other than transcription factors that regulate the rapid cell shape changes associated with chemotactic movement. Interestingly, Erk2 regulation of GtaC translocation occurs in response to multiple chemoattractants that promote different cell fates (i.e., foraging versus multicellular development) suggesting that GtaC function might be different in these responses or that other signaling mechanisms might also be important in regulating gene expression for these different cell fates. The cAMP regulation of GtaC translocation has been previously described as a developmental timer mechanism to coordinate changes in gene expression with the repeated cAMP signaling that occurs during aggregation (Cai et al., 2014). The role that GtaC might play in foraging remains to be assessed but GtaC function could potentially play a more widespread role in the cessation of cell growth in response to nutrient limitations that triggers both foraging and multicellular development. However, the increased movement of *gtaC^-^* cells to folate supports the idea that GtaC suppresses the expression of the folate receptor Far1, perhaps as a mechanism to promote multicellular development. An earlier study suggests that GtaC binds to many sites in the genome and regulates a multitude of genes encoding cAMP receptors, cell-cell adhesion proteins, and other proteins that play important roles in early development (Santhanam et al., 2015). The mechanisms of gene regulation by GtaC function appear to be complex, involving both induction and repression of genes, and it is not known if GtaC functions independently or in combination with other transcription factors. GtaC function in response to folate and cAMP signaling could potentially have both separate and overlapping gene regulation targets similar to the potential substrate targets of Erk2 phosphorylation in response to these chemoattractants (Nichols et al., 2019).

The analysis of Erk2 preferred phosphorylation sites on GtaC revealed that almost any combination of these sites would allow translocation of the transcription factor to the cytoplasm suggesting that the translocation process might be mediated by multiple changes in GtaC conformation. The precise mechanism of nuclear export is not known but sequences near the nuclear localization signal have been reported to resemble features of nuclear export signals in other systems. It is possible that exposure of a nuclear export signal could mediate the export but associations with other exported proteins might also allow for cytoplasmic localization. As previously reported, GtaC has sites that might be phosphorylated by other classes of protein kinases, such as glycogen synthase kinase (GSK) or cAMP-dependent protein kinase (PKA), but thus far only the loss of Erk2 preferred phosphorylation sites has severely impacted the translocation process. Gsk3 function in *Dictyostelium* has been previously shown to affect the efficiency of GFP-GtaC translocation in response to cAMP and GSKs typically recognize substrates already phosphorylated by other protein kinases, suggesting that phosphorylation of GtaC by Gsk3 might be a secondary modification (Cai et al., 2014). Whether or not the Erk2 phosphorylation sites impact the ability of GtaC to serve as a substrate for other protein kinases remains to be determined. The multiple shifts in electrophoretic gel migration suggest GFP-GtaC undergoes multiple phosphorylations or other modification after chemoattractant stimulation suggesting that GtaC regulation might be complex (Cai et al., 2014). The distribution of the Erk2 preferred or other protein kinase phosphorylation sites in the C-terminal half of GtaC does not easily explain the requirement of an N-terminal region (residues 180-222) for cytoplasmic translocation. Perhaps this N-terminal region contains binding sites necessary for protein kinase interactions or folds over near the C-terminus to regulate the exposure of signals that regulate GtaC translocation.

In mammalian cells, the atypical MAPK, MAPK15 (a.k.a. Erk8), has been associated with general cellular stress through protein binding analysis, heterologous expression, or RNA silencing but thus far no specific signals or signaling pathways have been identified that cause rapid activation of the MAPK like that observed for chemoattractant activation of Erk2 in *Dictyostelium* (Chia et al., 2014; Colecchia et al., 2018; Groehler and Lannigan, 2010; Hasygar and Hietakangas, 2014; Iavarone et al., 2006; Klevernic et al., 2009; Liwak-Muir et al., 2016; Rossi et al., 2011; Saelzler et al., 2006; Zacharogianni et al., 2011). If MAPK15 has roles in cell movement and development similar to that of Erk2 in *Dictyostelium* then disrupting the MAPK15 gene could potentially impact early embryonic development making the genetic analysis of MAPK15 in mammals challenging. Many important phenotypes associated with the loss of Erk2 in *Dictyostelium*, including defects in chemotaxis and transcription factor translocation, were not revealed with hypomorphic *erk2^-^* alleles suggesting that reduced levels of atypical MAPK expression, like those achieved through RNA silencing, might not be sufficient to uncover all MAPK associated phenotypes. However, developing a kinase translocation reporter for atypical MAPKs in mammals could provide a powerful tool for investigating the signals that activate MAPK15. In *Dictyostelium* the GFP tagged GtaC protein serves as the first known kinase translocation reporter specific for an atypical MAPK. This reporter will likely benefit future studies in the identification of other signals, perhaps other chemoattractants or morphogens, that activate atypical MAPK function.

## Methods

### Strains and culturing

The *erk2*^-^, *erk1*^-^, *erk1*^-^*erk2*^-^, *gα4*^-^, and *gα5^-^* and strains were created from the parental axenic strain KAx3 and a thymidine auxotrophic derivative JH10 as previously described (Hadwiger and Firtel, 1992; Hadwiger et al., 1996; Schwebs and Hadwiger, 2015; Schwebs et al., 2018). The far1-strain was created from KAx3 cells as previously described (Pan et al., 2016). The *gtaC^-^* strain was created from JH10 cells through a disruption of the *gtaC* locus with an insertion of the *thyA* gene resulting in developmental phenotypes as previously described for *gtaC^-^* cells (Cai et al., 2014). Expression of wild-type gene expression vectors, pFar1-Y (Far1) and pHC326 (GtaC) were capable of rescuing chemotaxis and developmental phenotypes associated with the gene disruptions. *Dictyostelium* strains were grown in axenic HL-5 medium (with thymidine supplement for JH10 strain) or on lawns of Klebsiella aerogenes as previously described (Dynes and Firtel, 1989; Watts and Ashworth, 1970). Vectors were transformed into *Dictyostelium* strains using electroporation as previously described (Dynes and Firtel, 1989). Multiple clones from all transformations were assessed for developmental morphogenesis or GFP-GtaC reporter translocation.

### GFP-GtaC translocation assay

Strains expressing the pHC329 GFP-GtaC^c-s^ or derivatives were grown in fresh axenic medium for several h prior to harvesting and plating on coverslips attached to the bottom side of 10 mm holes drilled into the 60 mm diameter petri dishes. After cells were allowed to attach to the coverslip for 10 min unattached cells were removed by 2-3 washes with developmental buffer (DB) composed of phosphate buffer (12mM NaH_2_PO_4_ adjusted to pH 6.1 with KOH) with the addition of 1 mM MgCl2 and 0.5 mM CaCl2. A final solution of 100 μl solution of DB was placed over the attached cells. Within 6 min of the final DB wash, the coverslip petridishes were mounted on 60x oil-immersion objective of an Olympus XI89 spinning disc confocal microscope. The software Cellsens was used to program time lapse exposures at 30 sec intervals for a duration of 4-8 min. Chemoattractant solutions (cAMP and folate) were added to the cells during the second time-lapse interval to a final concentation of 10 nM cAMP or 1 μM folate, unless otherwise noted. ImageJ software was used to create videos of the time lapse and to quantify the translocation of the reporter from the nucleus to the cytoplasm. The mean pixel intensity of the nucleus was divided by the mean pixel intensity of the cytoplasm. Measurements of the nuclear/cytoplasmic ratio in the stimulated cells were compared to an average of the ratios obtained from the first two time-lapse images representing unstimulated cells. The middle 50% values were used to generate the plotted data due to transient variability of values in some cells undergoing movement/shape changes that temporarily cause the nucleus to leave the focal plane.

### GFP-GtaC^c-s^ mobility shift assay

WT, *erk1*^-^, *erk2*^-^, and *erk2*^-^(Erk2) cells expressing GFP-GtaC^CS^ were starved in DB for 1 h. Cells were then washed with cold DB, resuspended in DB to a density of 2 × 10^7^ cells/ml, and kept on ice before assay. Cells were stimulated with 1 μM of cAMP at room temperature, lysed in SDS sample buffer at various time points before or after stimulation, and boiled for 5 min. Immunoblotting was performed as described before using an anti-GFP antibody (Roche, cat.#11814460001) (Kamimura et al., 2009).

### *In vitro* phosphorylation assay

To purify GtaC as the substrate, GFP-GtaC/*gtaC^-^* cells were washed with cold DB, lysed at a density of 1 × 10^7^ cells/ml by adding equal volume of 2× lysis buffer [50 mM Tris (pH 7.5), 200 mM NaCl, 1% NP-40, 100 mM NaF, 50 mM sodium pyrophosphate, 4 mM Na_3_VO_4_, 2x Complete EDTA-free protease inhibitor mixture], and incubated on ice for 5 min. Cleared lysate was incubated with GFP-Trap (ChromoTek) beads at 4°C for 1.5 hours. Beads were washed with lysis buffer and elution buffer [25 mM Tris (pH 7.5), 100 mM NaCl, 0.1% NP-40] and kept on ice before assay. To purify Erk1 and Erk2, Flag-Erk1/*erk1*^-^ and Flag-Erk2/*erk2*^-^ cells were collected before and after cAMP stimulation (1 μM of cAMP for 1 min), washed with DB, and lysed at a density of 2 × 10^7^ cells/ml by adding equal volume of 2 x lysis buffer. Cleared lysate was incubated with of anti-Flag M2 affinity resin (Sigma, cat. #F2426) at 4°C for 2 hours. Beads were washed with lysis buffer and elution buffer. Proteins were eluted by the addition of elution buffer containing 3 x Flag peptide (Sigma, cat. #F4799) at 4°C for 30 min. For kinase reaction, 10 μl of GFP-GtaC-containing beads were mixed with 25 μl of Flag-Erk1 or Flag-Erk2-containing eluate. Reaction was initiated by the addition of adenosine 5’-triphosphate (ATP) mix to a final concentration of 10 mM MgCl_2_, 5 mM DTT, and 0.3 mM ATP. Reaction was allowed to proceed for 15 min at room temperature and stopped by the addition of protein sample buffer.

### Vector construction

Expression vectors pHC326 (GFP-GtaC) and pHC329 (GFP-GtaC^c-s^, C500S/C503S) have been previously described (Cai et al., 2014). Erk2 phosphorylation site alterations at residues S357, S380, S386, and T492 were created through PCR mutagenesis of an intron-less version of pHC329 using the oligonucleotides (#1-14) and HiFi assembly (NEB) (Fig. S6). All of the phosphorylation site mutant vectors were further modified to restore the wild-type sequence in the zinc finger domain (S500C/S503C) using oligonucleotides (#15,16) (Fig. S6). All modified vectors were sequenced to verify changes in the sequence. The *gtaC::thyA* gene disruption was created through the insertion of a *thyA* gene fragment, amplified with oligonucleotides (# (o into a unique *Eco*RI site in the *gtaC* open reading frame and a *Bgl*II fragment containing the disruption site and flanking sequences was excised from a vector and electroporated into JH10 cells.

### Chemotaxis and developmental assays

Above agar chemotaxis assays to folate were conducted as previously described by placing droplets of cell suspensions on nonnutrient agar plates followed by the placement of droplets of 100 μM folate (approximately 2 mm away from the cell droplets (Nguyen et al., 2010). Images of cells acquired after plating and 3 h later were compared to measure the maximum distance traveled by the leading edge of cells. For cAMP assays cells were shaken in phosphate buffer for 3 h prior to plating on nonnutrient plates. The developmental morphogenesis of *gtaC^-^* cells, with or without GFP-GtaC vectors, was analyzed by transferring cells from clonal colonies in axenic culture onto a bacterial lawn. Images of *Dictyostelium* plaques on the lawn were acquired 2-3 days after the inoculation. Multiple transformants were analyzed for each strain.

## Author contributions

JAH and HC designed experiments, created strains and mutant alleles, and conducted most experiments. RGA and SH performed data analysis. All authors contributed to the manuscript writing and editing.

## Acknowledgement

This work was supported by the grants NIGMS R15 GM131269-01 and OCAST HR13-36 to JAH and National Science Foundation of China 32170701 to HC. The authors thank Alex Mason, Elise Ballinger, and Stormie Dreadfulwater for technical assistance.

## Supplementary Files

**Figure S1.** Videos of GFP-GtaC^c-s^ translocation in strains stimulated with 10 nM cAMP. Time-lapse video with images every 30 sec. Stimulant added at time of image 2 (contains red dot).

S1A: wild-type cells

S1B: *erk2*^-^ cells

S1C: *erk2*^-^(Erk2) cells

S1D: *erk1*^-^ cells

**Figure S2.** Videos of GFP-GtaC^c-s^ translocation in strains stimulated with 1 μM folate. Time-lapse video with images every 30 sec. Stimulant added at time of image 2 (contains red dot).

S2A: wild-type cells

S2B: *erk2*^-^ cells

S2C: *erk2*^-^(Erk2) cells

S2D: *erk1*^-^ cells

**Figure S3.**Videos of GFP-GtaC^c-s^ translocation in receptor and G protein mutant strains. Cells stimulated with 1 μM folate (S3A-C) or 10 nM cAMP (S3D-F). Time-lapse video with images every 30 sec. Stimulant added at time of image 2 (contains red dot).

S3A: *far1*^-^ cells

S3B: *gα4*^-^ cells

S3C: *gα5*^-^ cells

S3D: *far1*^-^ cells

S3E: *gα4*^-^ cells

S3F: *gα5*^-^ cells

**Figure S4.**
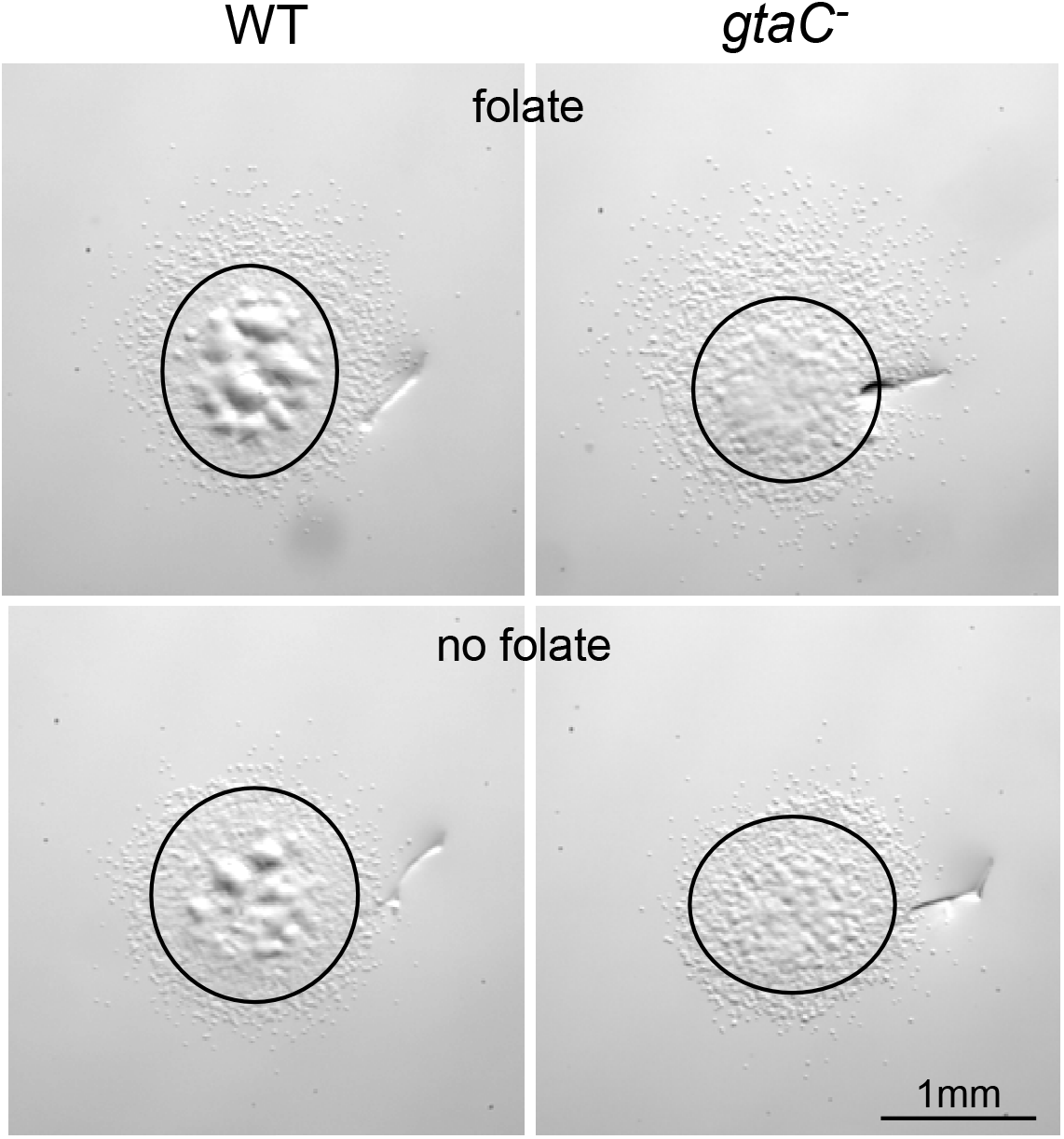
Images of wild-type or *gtaC^-^* cell chemotaxis to folate. Representative images of wild-type (WT) and *gtaC^-^* cells after 3 h in above agar chemotaxis assays to folate. In the appropriate panels the source of folate is above the cell droplet (upper side of panel). Circles indicate initial cell droplet perimeters and scars in the agar were used to align images before and after the assay. All images are the same magnification.

**Figure S5.**
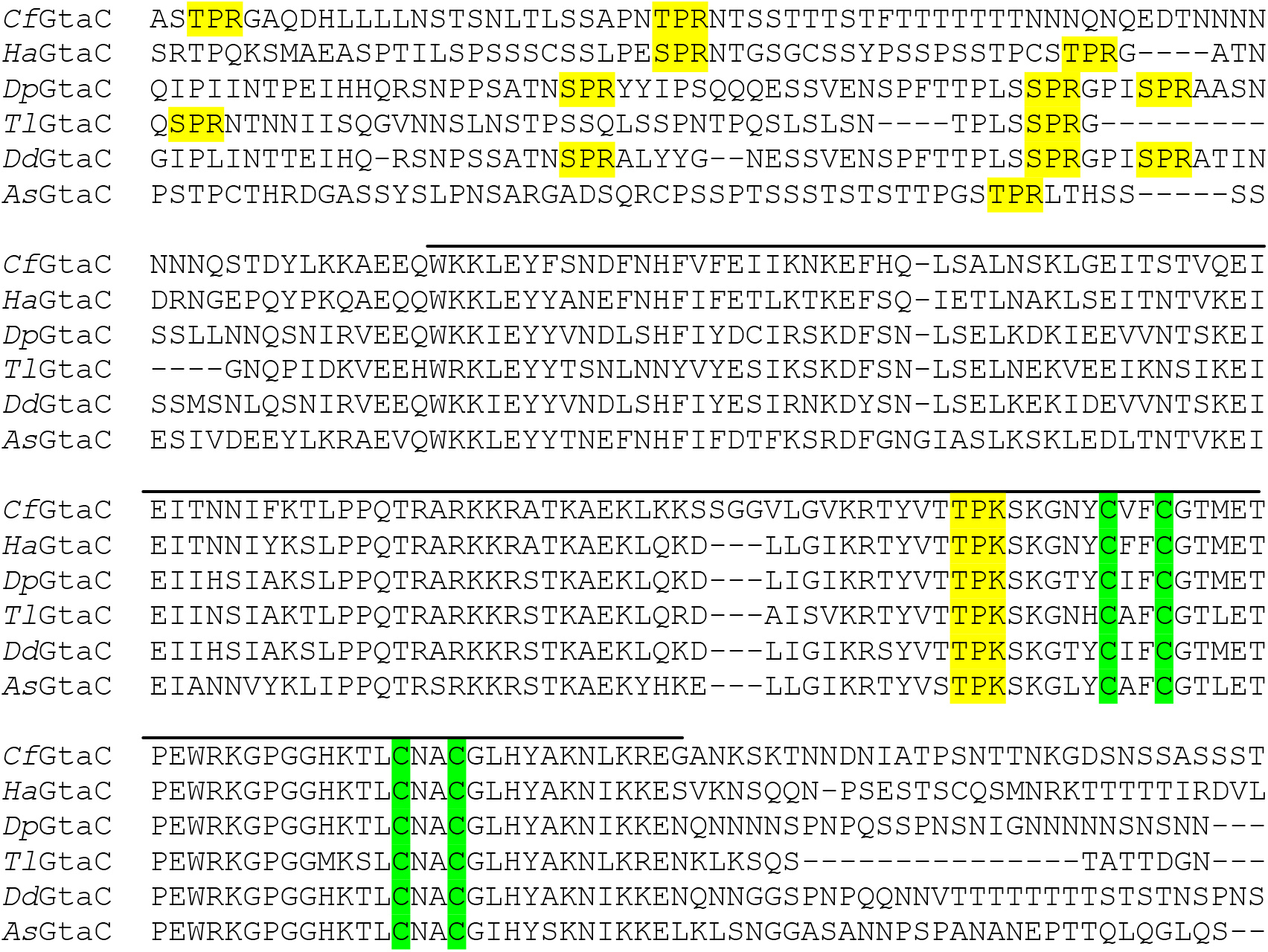
Alignment of Dictyostelid GtaC homologs. GtaC homologs from *Dictyostelium discoideium* (*Dd*) (residue 336),Cavenderia fasciculata(*Cf*) (residue 321), *Heterostelium album* (*Ha*) (residue 280),Dictyostelium purpureum (*Dp*) (residue 306), *Tieghemostelium lacteum* (*Tl*) (residue 226), and *Acytostelium subglobosum* (*As*)(residue 290) alignment using ClustalW and entire protein sequence (amino terminal portion not show prior to residue indicated). Erk2 preferred phosphorylation motifs (yellow), zinc finger conserved cysteines (green), and highly conserved region (line).

**Figure S6**. Videos of GFP-GtaC^c-s^ mutants with altered phosphorylation sites. Wild-type cells expressing translocation in folate strains stimulated with 1 μM folate. Timelapse video with images every 30 sec. Stimulant added at time of image 2 (contains red dot). S5A: GtaC^S357A^ (p911)

S5B: GtaC^T492A^ (p913)

S5C: GtaC^S380A^ (p926)

S5D: GtaC^S386A^ (p925)

S5E: GtaC^S357A,T492A^ (p937)

S5F: GtaC^S380A,S386A^ (p929)

S5G: GtaC^S357A,S380A,S386A^ (p931)

S5H: GtaC ^S380A,S386A,T492A^ (p932)

S5I: GtaC^S357A,S380A,S386A,T492A^ (p935)

**Figure S7.**
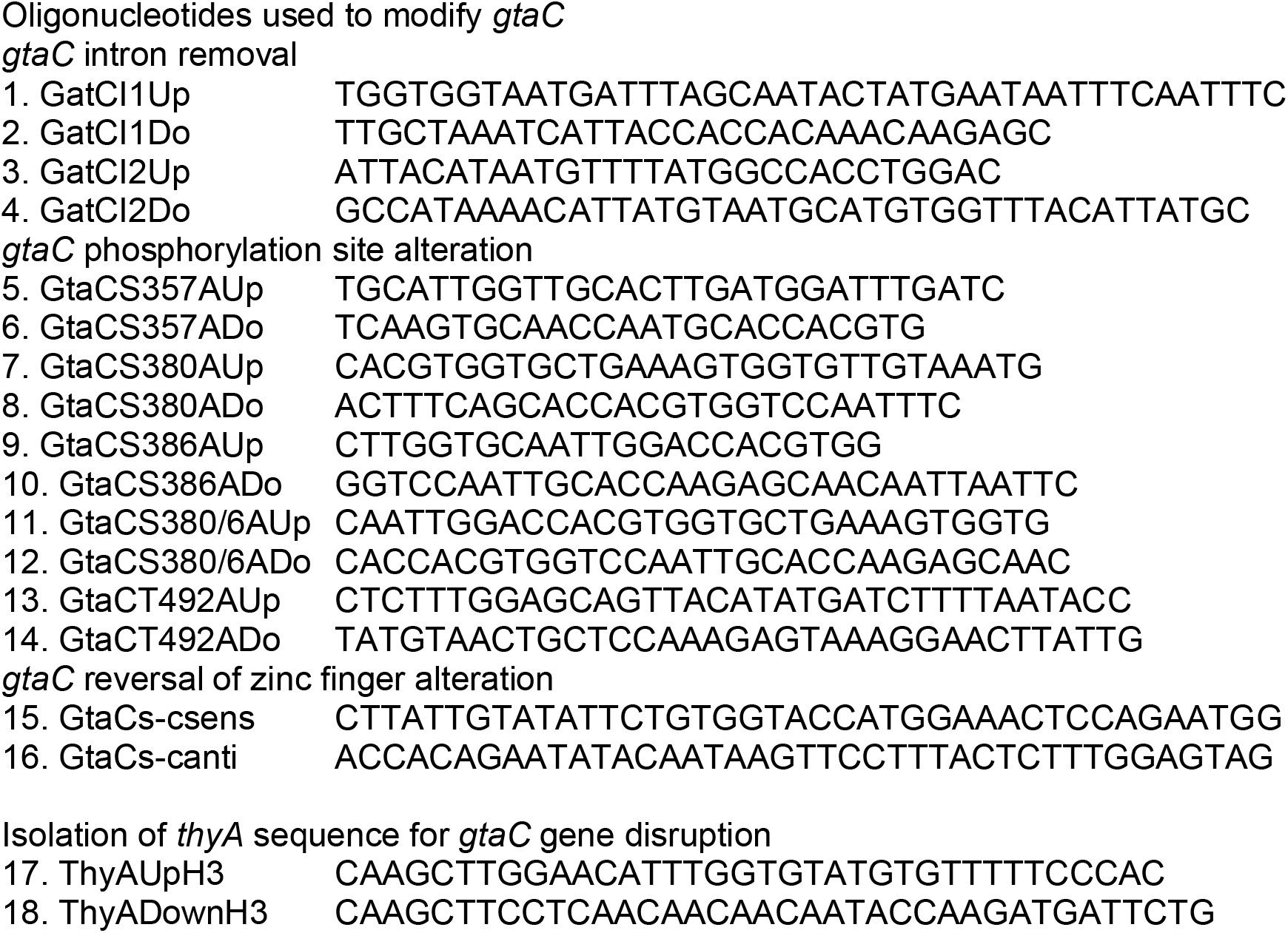
List of oligonucleotides. List of oligonucleotides used to modify *gtaC* (intron removal,alteration of phosphorylation sites, and reversal of zinc finger alteration) and isolate *thyA* sequence for gene disruption.

